# Environmental DNA reveals the structure of phytoplankton assemblages along a 2900-km transect in the Mississippi River

**DOI:** 10.1101/261727

**Authors:** Joseph M. Craine, Michael W. Henson, J. Cameron Thrash, Jordan Hanssen, Greg Spooner, Patrick Fleming, Markus Pukonen, Frederick Stahr, Sarah Spaulding, Noah Fierer

**Affiliations:** Jonah Ventures, 1600 Range St. #201, Boulder, CO, 80301 USA.; Department of Biological Sciences, Louisiana State University, Baton Rouge, LA, USA.; O.A.R. Northwest, Seattle, WA 98103, U.S.A.; School of Oceanography, University of Washington, Seattle, WA 98195, USA.; Institute of Arctic and Alpine Research, University of Colorado, Boulder, 80309 USA; Department of Ecology and Evolutionary Biology, University of Colorado, Boulder, CO80309, USA; Cooperative Institute for Research in Environmental Science, University of Colorado, Boulder, CO 80309

## Abstract

The environmental health of aquatic ecosystems is critical to society, yet traditional assessments of water quality have limited utility for some bodies of water such as large rivers. Sequencing of environmental DNA (eDNA) has the potential to complement if not replace traditional sampling of biotic assemblages for the purposes of reconstructing aquatic assemblages and, by proxy, assessing water quality. Despite this potential, there has been little testing of the ability of eDNA to reconstruct assemblages and their absolute and relative utility to infer water quality metrics. Here, we reconstruct phytoplankton communities by amplifying and sequencing DNA from a portion of the 23S rRNA region from filtered water samples along a 2900-km transect in the Mississippi River. Across the entire length, diatoms dominated the assemblage (72.6%) followed by cryptophytes (8.7%) and cyanobacteria (7.0%). There were no general trends in the abundances of these major taxa along the length of the river, but individual taxon abundance peaked in different regions. For example, the abundance of taxa genetically similar to *Melosira tropica* peaked at approximately 60% of all reads 2750 km upstream from the Gulf of Mexico, while taxa similar to *Skeletonema marinoi* began to increase below the confluence with the Missouri River until it reached approximately 30% of the reads at the Gulf of Mexico. There were four main clusters of samples based on phytoplankton abundance, two above the confluence with the Missouri and two below. Phytoplankton abundance was a poor predictor of NH_4_ ^+^ concentrations in the water, but predicted 61% and 80% of the variation in observed NO_3_ ^-^and PO_4_ ^3-^concentrations, respectively. Phytoplankton richness increased with increasing distance along the river, but was best explained by phosphate concentrations and water clarity. Along the Mississippi transect, there was similar structure to phytoplankton and bacterial assemblages, indicating that the two sets of organisms are responding to similar environmental factors. In all, the research here demonstrates the potential utility of metabarcoding for reconstructing aquatic assemblages, which might aid in conducting water quality assessments.

## Introduction

Inland freshwater systems provide vital services of drinking water, habitat for fisheries, irrigation for agriculture and recreation (Davies and Jackson 2006, American Sportfishing Association 2015). Yet, the ecological status of lakes, rivers, streams, and reservoirs is increasingly threatened by agriculture, roads, industry, mining, human waste, urbanization, and deforestation (Malmqvist and Rundle 2002, US Environmental Protection Agency 2015) Effective monitoring of water quality and the causes of water quality impairment are critical steps to maintaining freshwater resources, preventing further degradation, and guiding restoration efforts. Quantifying the state and dynamics of aquatic ecosystems is often best done indirectly by quantifying the structure of aquatic assemblages (Palaniappan et al. 2010, Young and Loomis 2014). Because each organism has a unique set of ecological traits and responds uniquely to environmental conditions, their abundance in waters is an indicator of environmental conditions such as salinity, temperature, oxygen levels, nutrient supplies, and turbidity (Karr 1999, Schoolmaster et al. 2012).

Fish and aquatic invertebrates are the two of the most common indicators quantified for the purpose of inferring water quality (Barbour et al. 1999, Stein et al. 2014a). Yet, assessments of these assemblages are currently labor intensive, slow, expensive, and often imprecise. For example, manually sampling fish or aquatic invertebrate communities can cost approximately US$2500 for a single site, limiting the number of sites that can be sampled (Stein et al. 2014a). Biotic assessments also are less effective for certain types of aquatic ecosystems. For example, even without fiscal constraints, assessments of fish and aquatic insect assemblages of large rivers can be exceedingly difficult. The efficiency of sampling fish in large rivers with traditional electrofishing is low (and seasonally variable) due to a number of factors such as turbidity (Goffaux et al. 2005, Reyjol et al. 2005, Lyon et al. 2014). Standard techniques for kicknetting insects or collecting exuviae do not work on large rivers (Buss et al. 2015).

One response to the constraints on sampling fish and insects for large rivers is to rely on other organisms such as diatoms for water quality assessments (Kelly and Whitton 1995, Stein et al. 2014a). Yet, traditional techniques for visually assessing the relative abundance of taxa such as diatoms is still expensive, subject to taxonomic bias, and constrained by low taxonomic resolution (Zimmermann et al. 2015). Given these constraints, next generation sequencing of environmental DNA has the potential to quantify assemblages of not only fish and aquatic insects, but also smaller organisms such as phytoplankton or bacteria (Mächler et al. 2014, Stein et al. 2014b, Barnes and Turner 2015, Thomsen and Willerslev 2015). To accomplish this, DNA present in the water is filtered and then regions in the genome are amplified and sequenced, providing information on the presence, if not relative abundance, of organisms. Depending on the regions of the genome amplified, different taxonomic groups can be sequenced, including bacteria, phytoplankton, arthropods, fish, and mammals (Jackson et al. 2014, Stein et al. 2014b, Cannon et al. 2016, Deiner et al. 2016, Olds et al. 2016). This potential is coupled with the ability to provide data for less cost, or improved taxonomic specificity, and at a faster rate. For example, water can be filtered and analyzed for environmental DNA at less than a tenth of the cost of traditional biotic assessments.

Despite this potential, there are few examples of successful application of metabarcoding for reconstructing phytoplankton assemblages and we have not started in earnest to assess whether these reconstructions have value on their own, no less relative to reconstructions generated with organisms (Hamsher et al. 2013). To better understand the potential of environmental DNA to reconstruct phytoplankton assemblages in a large river, we amplified and sequenced DNA using a 23S rRNA gene region primer pair (Sherwood and Presting 2007) specific to phytoplankton (hereafter, 23S) for 39 sites along over 2900 km of the Mississippi River in addition to its headwaters at Lake Itasca. The Mississippi River is one of the Great Rivers of the US and has been the subject of a number of studies attempting to assess the ecological health of its waters with biological assessment (Angradi et al. 2009, Kireta et al. 2012b, Bellinger et al. 2013). With these data, we examined the patterns of phytoplankton assemblages along the length of the river to determine how the structure and richness of these assemblages changed along the length of the river. As a first test, we compared relationships between the abundance of 23S OTUs and nutrient concentrations in the water. Next, to assess whether the factors structuring phytoplankton were similar to those structuring bacterial assemblages, we compared assemblage structure of 23S and bacterial 16S rRNA gene (hereafter, 16S) OTUs from a previous set of analyses (Henson et al. in review). This was followed with a comparison of the explanatory power of 23S and 16S OTUs to predict nutrient concentrations.

## Methods

### Sample acquisition

Duplicate water samples were collected from 39 sites along the Mississippi River from September 18, 2014 to November 26, 2014 (Henson et al. in review). The core of these sites spanned 2917 km, from Minneapolis, MN to the Gulf of Mexico. An additional sample was acquired from Lake Itasca, the headwaters of the river. At each site, 120 mL of water was filtered through a 2.7 μm GF/D filter (Whatman GE, New Jersey, USA) and then a 0.2 μm Sterivex filter (EMD Millipore, Darmstadt, Germany) with a sterile 60 mL syringe (BD, New Jersey, USA). The first 60 mL of flow-through water was collected and saved in an autoclaved, acid-washed 60 mL polycarbonate bottle. Filters and filtrate were stored on ice until they could be shipped to the laboratory for analyses. At each site, light penetration was assessed with a secchi disk (Wildco, Yulee, FL).

### Sample processing

DNA was extracted from filters with a MoBio PowerWater DNA kit (MoBio Laboratories, Carlsbad, CA) following the manufacturer’s protocol. If there was sufficient DNA remaining from previous analyses (Henson et al. in review), DNA from the two fractions for a site were combined. For some sites, DNA from the two fractions taken from the two replicate samples were combined. Phytoplankton sequences were amplified at the 23S rRNA gene region, which is located on the chloroplast and can amplify DNA from taxa such as cyanobacteria, green algae, and diatoms (Sherwood and Presting, 2007). Initial PCR amplification included Promega Mastermix, forward and reverse primers, gDNA, and DNase/RNase-free H2O. After an initial 3-minute period at 94°C, DNA was PCR amplified for 40 cycles at 94°C (30 seconds), 55°C (45 seconds) and 72°C (60 seconds), followed by 10 minutes at 72°C. Products were then visualized on an 2% agarose gel. 20μl of the PCR amplicon was used for PCR clean-up using ExoI/SAP reaction. In order to index the amplicons with a unique identifier sequence, the first PCR stage was followed by an indexing 8-cycle PCR reaction to attach 10-bp error-correcting barcodes unique to each sample to the pooled amplicons. These products were again visualized on a 2% agarose gel and checked for band intensity and that amplicons are the correct size. PCR products were purified and normalized using the Life Technologies SequalPrep Normalization kit and samples pooled together. Amplicons were sequenced on an Illumina MiSeq at the University of Colorado Boulder BioFrontiers Sequencing Center running the v2 500-cycle kit.

For nutrient analyses, filtrate was previously analyzed colorimetrically for [NH_4_^+^], [NO_3_^-^], and [PO_4_^3-^] at the University of Washington Marine Chemistry Laboratory as described in Henson et al. (in review).

### Bioinformatic processing

Sequences were demulitplexed using a python script. Paired end reads were then merged using fastq_merge pairs. Since merged reads often extended beyond the amplicon region of the sequencing construct, we used fastx_clipper to trim primer and adaptor regions from both ends (https://github.com/agordon/fastx_toolkit). Sequences lacking a primer region on both ends of the merged reads were discarded. Sequences were quality trimmed to have a maximum expected number of errors per read of less than 0.1 and only sequences with more than 3 identical replicates were included in downstream analyses. BLASTN 2.2.30+ was run locally, with a representative sequence for each OTU as the query and the current NCBI nt nucleotide and taxonomy database as the reference. The tabular BLAST hit tables for each OTU representative were then parsed so only hits with > 97% query coverage and identity were kept.

Sequences were clustered into OTUs at the ≥ 97% sequence similarity level and sequence abundance counts for each OTU were determined using the usearch7 approach. The National Center for Biotechnology Information (NCBI) genus names associated with each hit were used to populate the OTU taxonomy assignment lists. Sequences that did not match over 90% of the query length and did not have at least 85% identity were considered unclassified. Otherwise the top BLASTn hit was used.

### Statistical analyses

To quantify the accumulation of 23S OTUs with increasing numbers of samples, we used the *specaccum* function of the *vegan* package with the Lomolino function to describe the curves (Oksanen et al. 2017).

Hierarchical clustering of 23S was based on Ward’s minimum variance method. A heat map was generated with the *heatmap.2* function of the *gplots* package (Warnes et al. 2016) using distance matrices created from the relative abundance of the top 50 23S OTUs. To identify taxa disproportionately associated with the 8 major clusters, indicator values were calculated for each of the top 50 OTUs based on abundance of occurrence (Dufrêne and Legendre 1997).

To assess the relationships between nutrient concentrations and 23S OTU abundance, forward stepwise regression was performed for [NH_4_^+^], [NO_3_^-^], and [PO_4_^3-^] with the top 50 23S OTUs (*P* < 0.01 for entry). To assess the relationships between phytoplankton richness and predictors, all singletons were removed from the abundance of reads, phytoplankton richness was first rarefied to the minimum number of reads for the sample set (4,212) and then regressed in a backwards elimination stepwise regression with nutrient concentration data, distance along the river, and secchi disk depth.

To compare 23S and 16S patterns, we restricted 16S data to the top 100 OTUs, representing 76% of the total reads. Previously, the 16S region was sequenced for the two particle size fractions independently (Henson et al. in review). Here, bacterial OTU abundance was averaged for the two fractions for a given sample. Mantel tests ( *mantel* function of the *vegan* package) assessed Pearson correlations among assemblage similarity matrices, which were based on Euclidean distances. A cophenetic correlation was assessed for the 23S and 16S distance matrices using the *cor_cophenetic* function of *dendextend* package (Galili 2015). To visualize similarity in clustering between 23S and 16S OTU abundances, a tanglegram was generated using the *tanglegram* function of *dendextend* package based on the 23S hierarchical clustering and a new hierarchical clustering of 16S data also based on Ward’s minimum variance method. The same stepwise regression technique on nutrient concentrations was used for the top 50 16S OTUs as was done for the 23S OTUs.

All statistical analyses were conducted in R 3.2.5 using Rstudio v. 1.0.136 except the stepwise regressions, which were computed in JMP v. 13.0.0 (SAS Institute, Cary NC, USA).

## Results

Across all samples, the most abundant phytoplankton OTU was for taxa similar to *Thalassiosira rotula,* which represented 37.6% of all reads. The next most abundant OTU was for taxa similar to the diatom *Melosira tropica,* which represented 15.8% of all reads. In general, the top 10 OTUs represented 80.9% of all reads and the top 50 OTUs represented 96.4% of all reads. Among the top 50 OTUs, 72.6% of the reads were from Bacillariophyta, 8.7% were from Cryptophyta, and 7.0% were from Cyanobacteria. Chlorophyta and Eustigmatophyceae comprised 3.5% and 3.2% of the reads, respectively. Examining the pattern of OUT accumulation, OTU abundance is predicted to asymptote at 447 OTUs with half of this occurring in 11.8 samples (Figure 1). Mean richness after rarefication was 55.3 ± 15.9 (s.d.) OTUs per sample. Mean richness increased at a rate of 7.1 ± 1.7 species per 1000 km (r^2^ = 0.23, *P* < 0.001).

**Figure 1.**
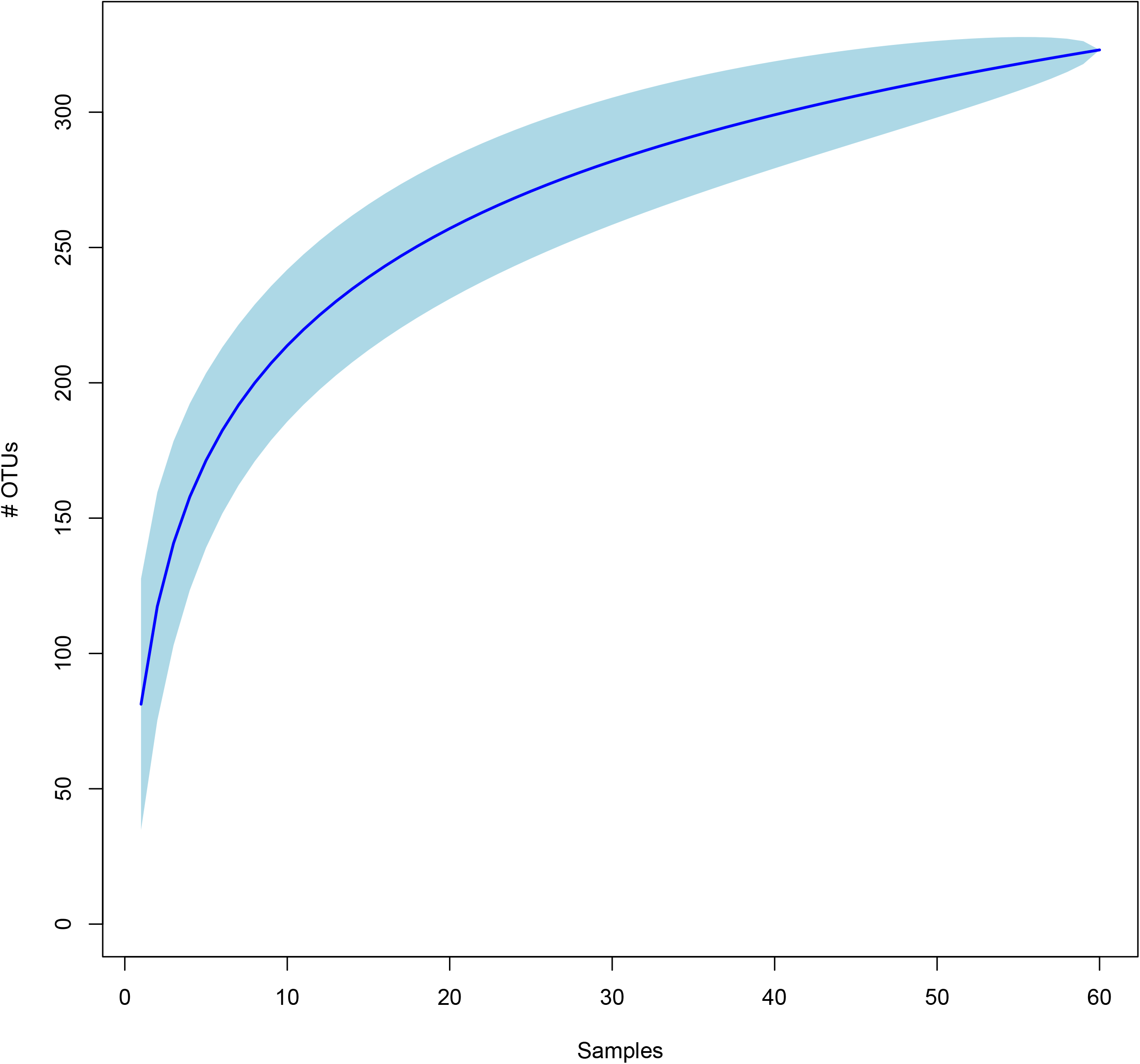
Accumulation of OTUs with additional samples. 50% of the accumulation of OTUs occurs with 11.8 samples and richness is predicted to asymptote at 447 samples.

Phytoplankton had different patterns of distribution along the length of the river (Figure 2). Among the four most abundant OTUs, *Melosira tropica* OTU abundance peaked at approximately 60% of all reads 2750 km upstream from the Gulf of Mexico, while *Thalassiosira rotula* OTU abundance peaked at approximately 90% of all reads approximately 2250 km from the Gulf. In contrast, *Cyclotella* sp. WC03 (OTU 48) did not peak until ~1300 km from the Gulf (17% of all reads) and the *Skeletonema marinoi* OTU continued to increase below the confluence with the Missouri River, until it reached approximately 30% of the reads at the Gulf of Mexico. There were no general trends in the abundance of phytoplankton groups with respect to distance along the river when read abundance for the top 50 OTUs was aggregated by phylum (Figure 3).

**Figure 2.**
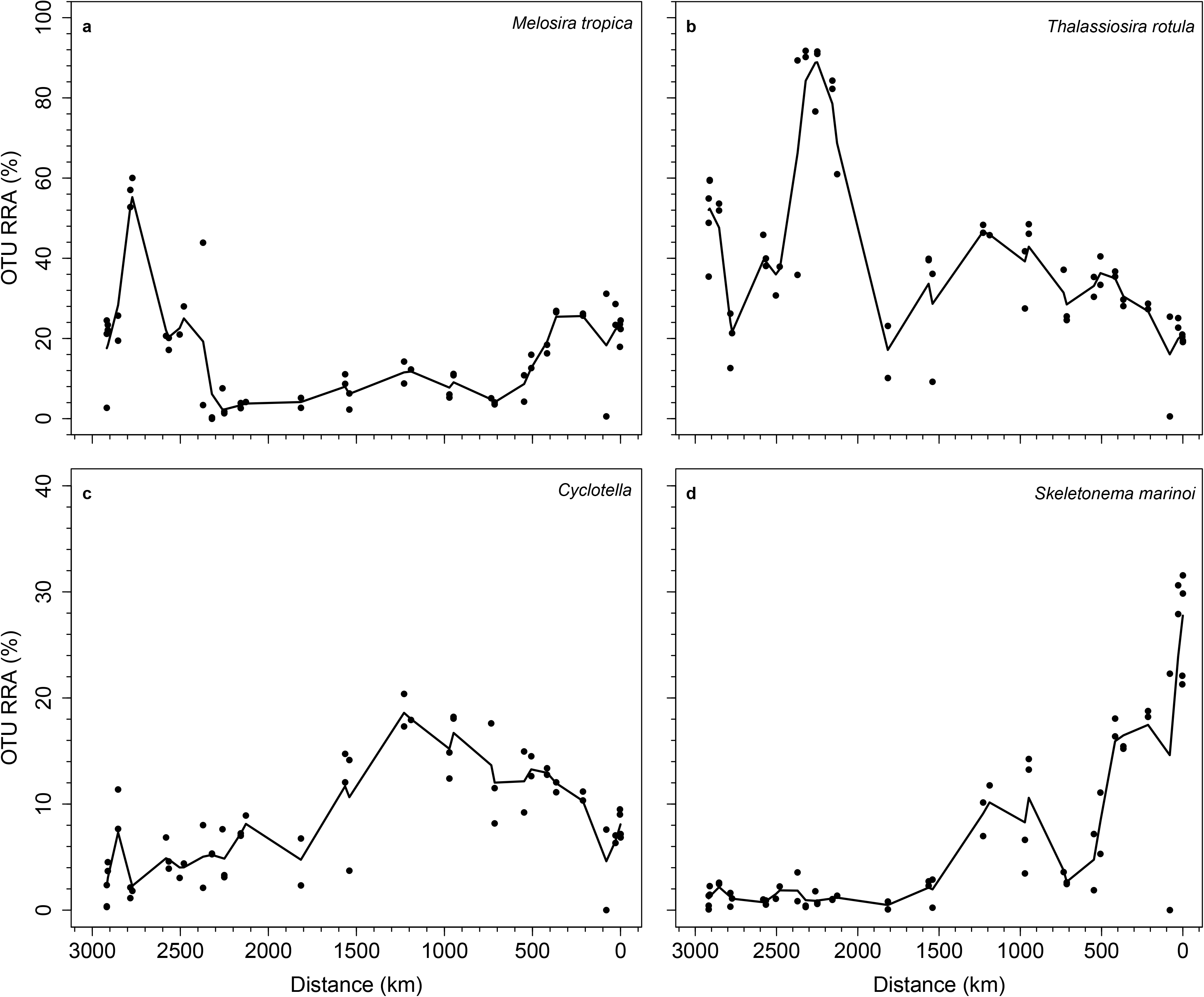
Relative read abundance of four most abundant OTUs as a function of distance from the mouth of the Mississippi River.

**Figure 3.**
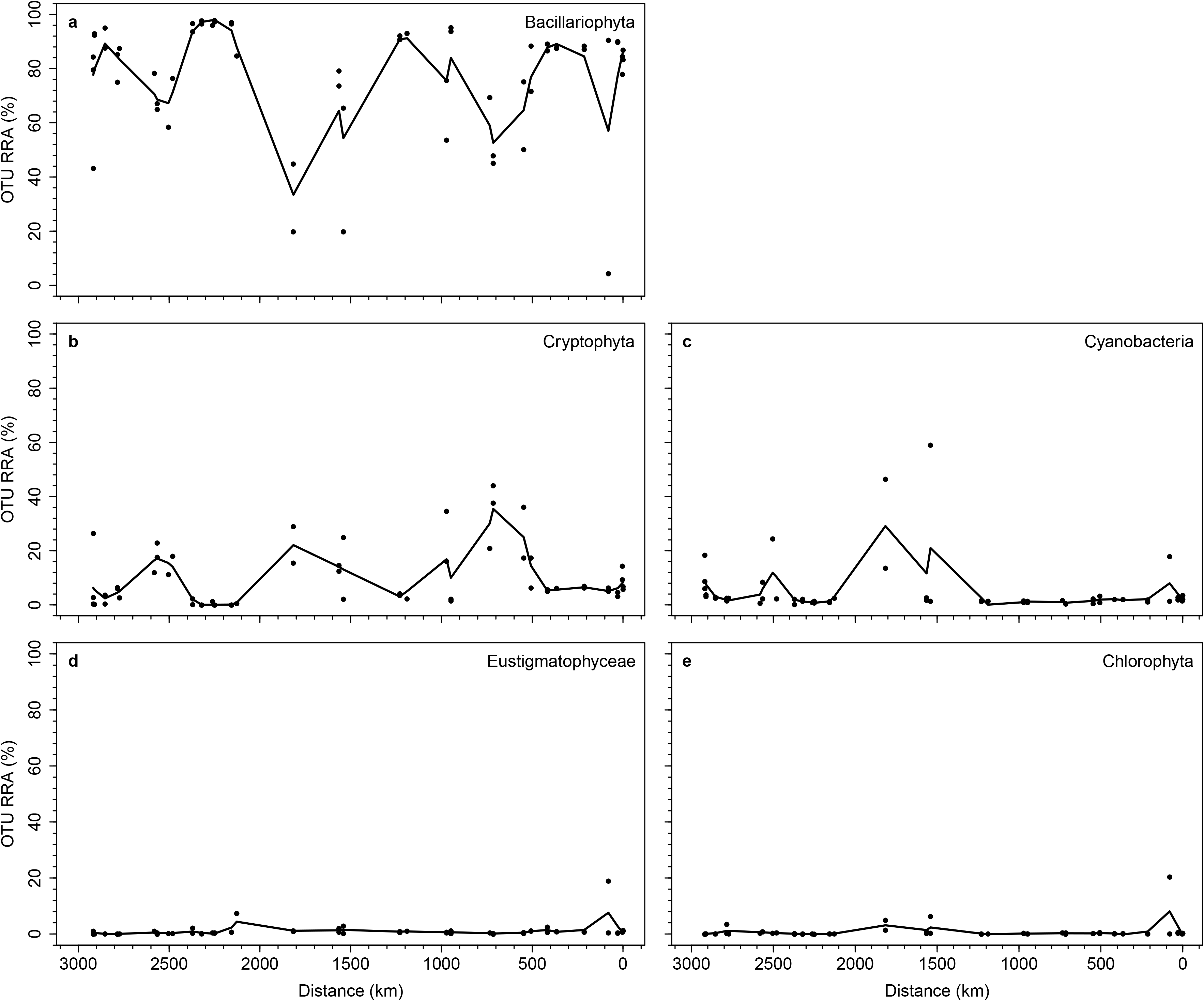
Relative read abundance of five main taxonomic groups as a function of distance from the mouth of the Mississippi River.

### Clustering of sites

The phytoplankton of Lake Itasca was the most unique set of OTUs and did not cluster with any other samples (Figure 4). The Lake Itasca assemblage was characterized by the abundance of chrysophyte species similar to *Ochromonas danica*, dinoflagellates similar to *Dinophysis fortii* and species similar to the yellow-green alga *Trachydiscus minutus* (Table 1). Beyond Lake Itasca, four other main clusters of sites were identified, which encompassed 57 of the remaining 61 samples. The first cluster contained 17 of the 27 samples taken upstream of the confluence with the Missouri River (Figure 4). These samples were indicated by their abundances of taxa similar in sequence to *Thalassiosira rotula* ( *P* = 0.003; Table 1). The second cluster consisted of 8 samples in the Upper Mississippi that ranged along 300 km from Lake Pepin in Minnesota to Dubuque, Iowa. These sites were indicated by their abundances of species similar in sequence to the dinoflagellate *Gymnodinium eucyaneum*, the diatom *Tenuicylindrus* sp., the cryptomonad *Cryptochloris*, and the diatom *Melosira tropica* (Table 1). The third main cluster denoted 20 of the samples below the confluence with the Missouri River, primarily by their abundance of taxa similar in sequence to *Skeletonema marinoi*. The fourth main cluster contained 12 samples below the Missouri River confluence from above Vicksburg, MS to just below Three Rivers Wildlife Management Area. These samples were indicated by their abundances of species similar in sequence to the cryptomonads *Teleaulax acuta*, *Cryptomonas* sp., and *Plagioselmis nannoplanctica* as well as two diatom OTUs for species similar in sequence to *Cyclotella* sp. (Table 1).

**Figure 4.**
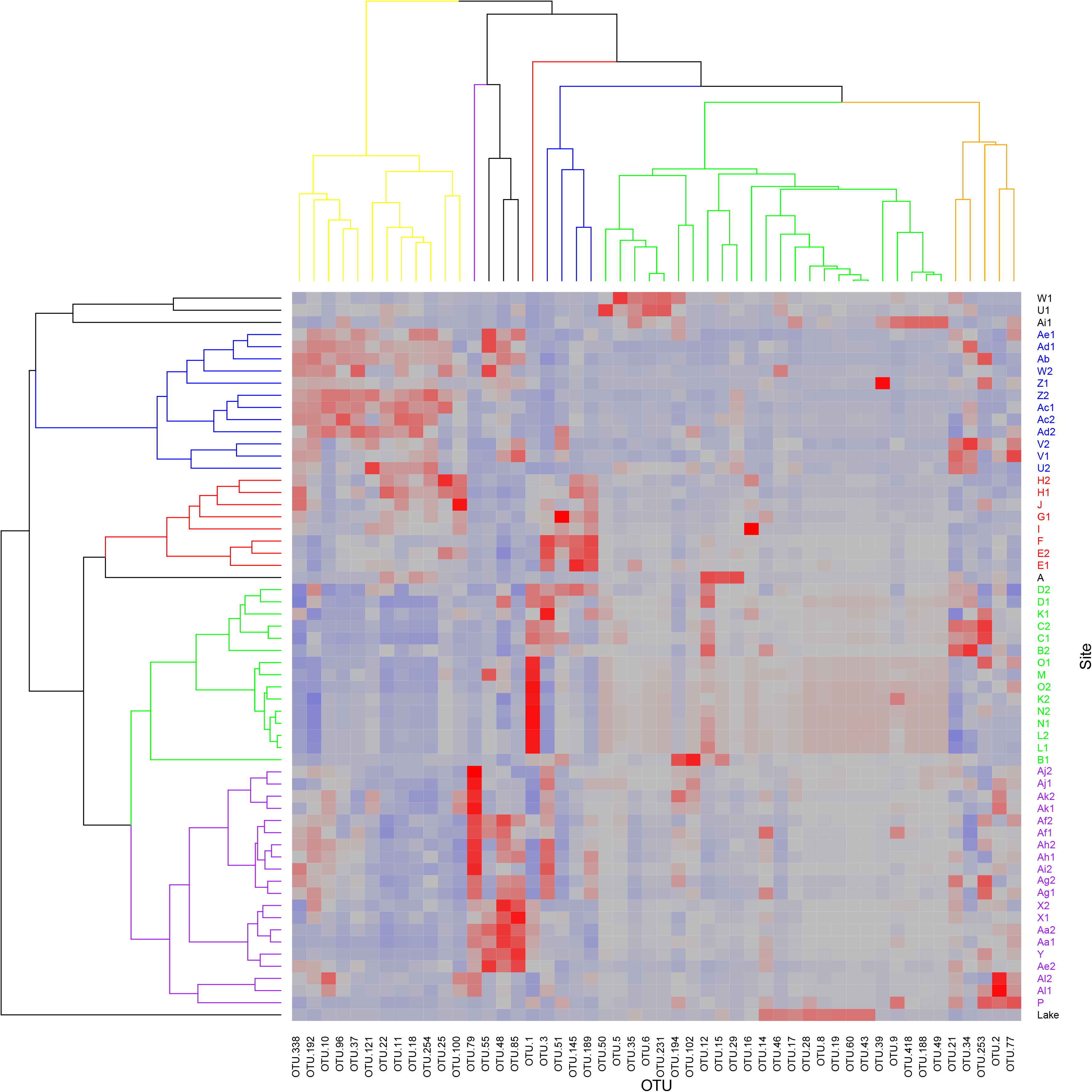
Heat map of abundances of OTUs at sites along the Mississippi River based on the standardized relative abundance of the 50 most abundant 23S OTUs. Blue indicates a low relative abundance and red high with gray intermediate. Sites and OTUs were clustered hierarchically based on dissimilarity index of relative abundances. Four major site clusters shown in color including Cluster 2 (purple), Cluster 5(green), Cluster 6 (red), and Cluster 3 (blue).

**Table 1.**
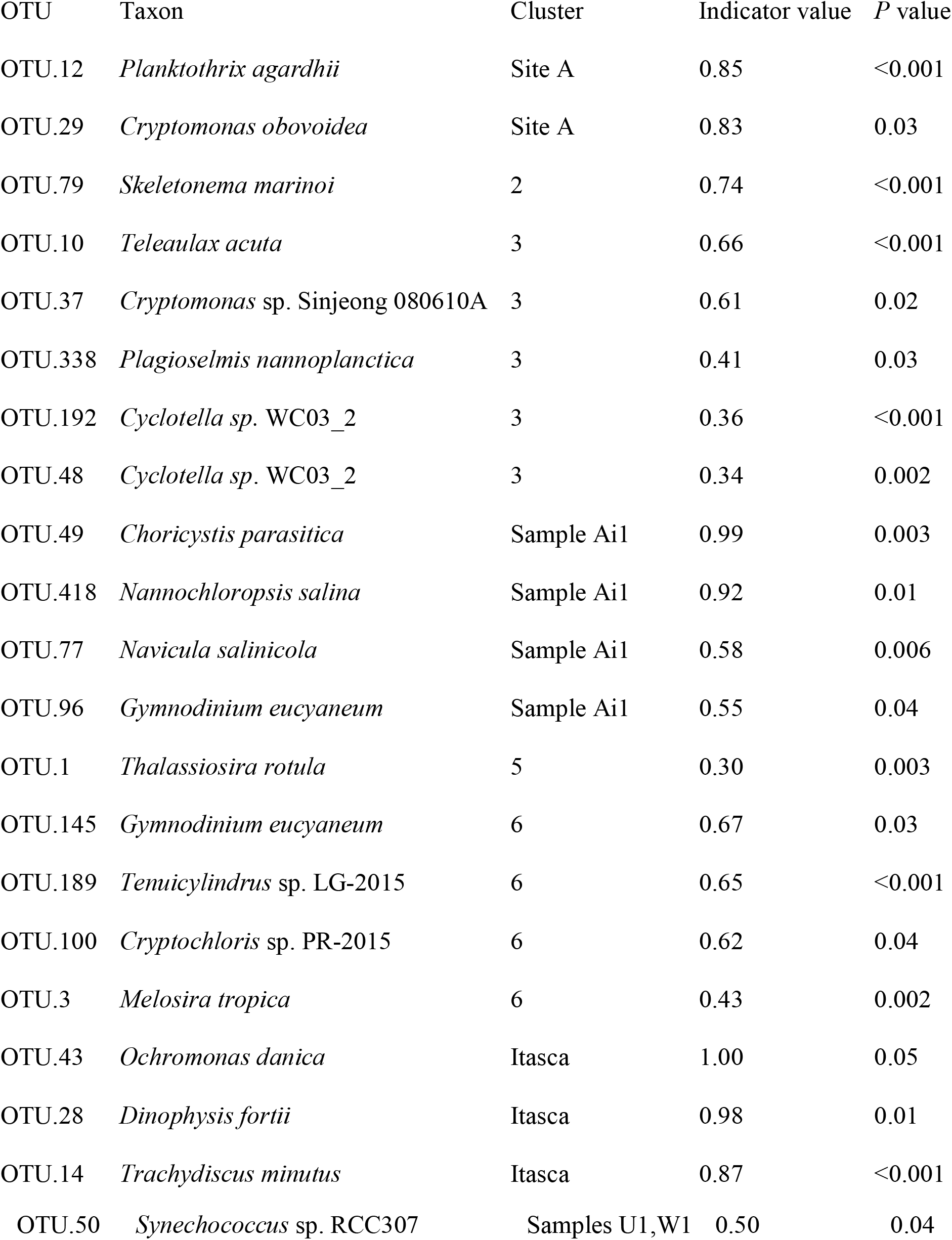
Table of indicator values for different clusters. See Figure 4 for which sites belong to each cluster.

### Relationships with nutrient data

The best predictor of NH_4_ ^+^ concentrations at a given location was the abundance of diatoms similar in sequence to *Sellaphora pupula*, which explained 38% of variation in [NH_4_^+^] (Table 2), but mostly as a result of two sites having high [NH_4_^+^] (> 25 μg L^-1^) and abundances of *Sellaphora pupula*. After this OTU, the abundances of no other phytoplankton OTU predicted NH_4_ ^+^ concentrations ( *P >* 0.01 for all OTUs). [NO_3_^-^] was best predicted by 4 diatom OTUs, which explained 61% of the variation in [NO_3_^-^]. Nitrate concentrations decreased with increasing abundances of species similar in sequence to *Melosira tropica* and increased with increasing abundances of species similar in sequence to *Cyclotella* sp. WC03, *Navicula salinicola,* and *Dinophysis fortii* (Table 2). 80% of the variation in [PO_4_^3-^] was explained by the abundances of six diatom OTUs (Table 2). [PO_4_^3-^] increased with increasing abundances of species similar in sequence to *Skeletonema marinoi*, *Cyanobium* sp. *Navicula salinicola*, *Cyclotella* sp. WC03, *Cryptomonas ovata*, and *Dinophysis fortii.* Phytoplankton OTU richness increased downstream ( *P* < 0.01), but in the backwards elimination stepwise regression, phytoplankton OTU richness (intercept = 16.42 ± 6.83; *P* = 0.02) increased with increasing secchi disk depth (0.295 ± 0.071 OTUs cm^-1^; *P* < 0.001) and with increasing [PO_4_^3-^] (0.328 ± 0.048 OTUs (μg L ); *P* < 0.001) (Figure 5).

**Figure 5.**
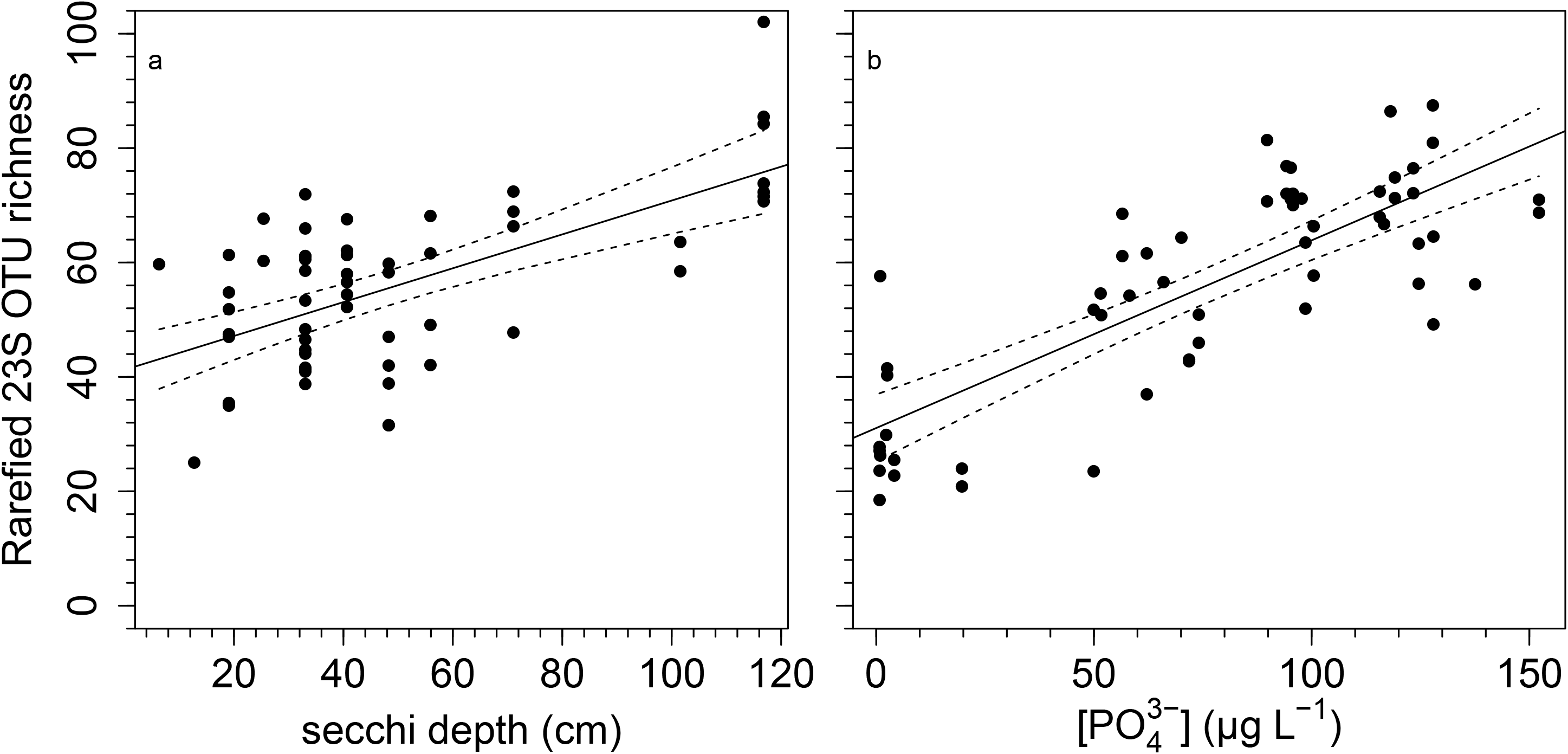
Partial residual plots of rarefied OTU richness as a function of (a) secchi disk depth and (b) [PO_4_^3-^]. Non-significant variables include distance down the river, [NO_3_^-^], and [NH_4_^+^].

**Table 2.** 23S OTU predictors of nutrient concentrations including sums of squares, estimates (μg L^-1^), and *P* values. Coefficients of variation for [NH_4_^+^], [NO_3_^-^], and [PO_4_ ^3-^] were 0.38, 0.61, 0.80, respectively.

| Nutrient | Variable | %SS | Estimate | $P$ value |
| --- | --- | --- | --- | --- |
| $[\text{NH}_4^+]$ | Intercept | | $3.2 \pm 0.8$ | <0.001 |
| | *Sellaphora pupula* | 100% | $492.4 \pm 75.7$ | <0.001 |
| $[\text{NO}_3^-]$ | Intercept | | $1046 \pm 148.4$ | <0.001 |
| | *Cyclotella* sp. WC03_2 | 34.1% | $-1947.5 \pm 453.1$ | <0.001 |
| | *Melosira tropica* | 23.0% | $4152.7 \pm 1177.9$ | <0.001 |
| | *Navicula salinicola* | 22.6% | $67288.8 \pm 19245.2$ | <0.001 |
| | *Dinophysis fortii* | 20.3% | $5803301 \pm 1749619.8$ | 0.002 |
| $[\text{PO}_4^{3-}]$ | Intercept | | $2.4 \pm 6.4$ | 0.7095 |
| | *Cyclotella* sp. WC03_2 | 8.2% | $248.3 \pm 61.1$ | <0.001 |
| | *Skeletonema marinoi* | 38.2% | $317.3 \pm 36.2$ | <0.001 |
| | *Cyanobium* sp. PCC 7009 | 27.6% | $352.2 \pm 47.2$ | <0.001 |
| | *Cryptomonas ovata* | 9.9% | $1597.8 \pm 357.6$ | <0.001 |
| | *Navicula salinicola* | 9.9% | $3977.9 \pm 891.1$ | <0.001 |
| | *Dinophysis fortii* | 6.2% | $299742.4 \pm 84789.6$ | <0.001 |

### Comparing phytoplankton and bacteria

Comparing distance matrices with a Mantel test, 23S and 16S assemblages were correlated ( *r* =0.44, *P* < 0.001). Similarly, the hierarchical clustering of sites based on 23S and 16S assemblages were correlated (cophenetic correlation, *r* = 0.43), revealing structural similarity in the two assemblages. For example, comparing the dendrograms, paired samples often clustered together for both the 23S and 16S assemblages, such as the D samples and Aa samples. Also, sites U and W were more similar to one another than other sites for both 23S and 16S (Figure 6). Stepping back to the broader patterns, the major clusters of sites in the 23S data were also largely present for the 16S data, though the relative positions within this cluster were mixed. Some differences in the clustering between the two sets of samples were likely due to stochasticity or contamination in individual samples for one primer pair. For example, with the 23S data, site P clustered with the Al sites. Yet, in the 16S data it clustered more closely with the adjacent O sites.

**Figure 6.**
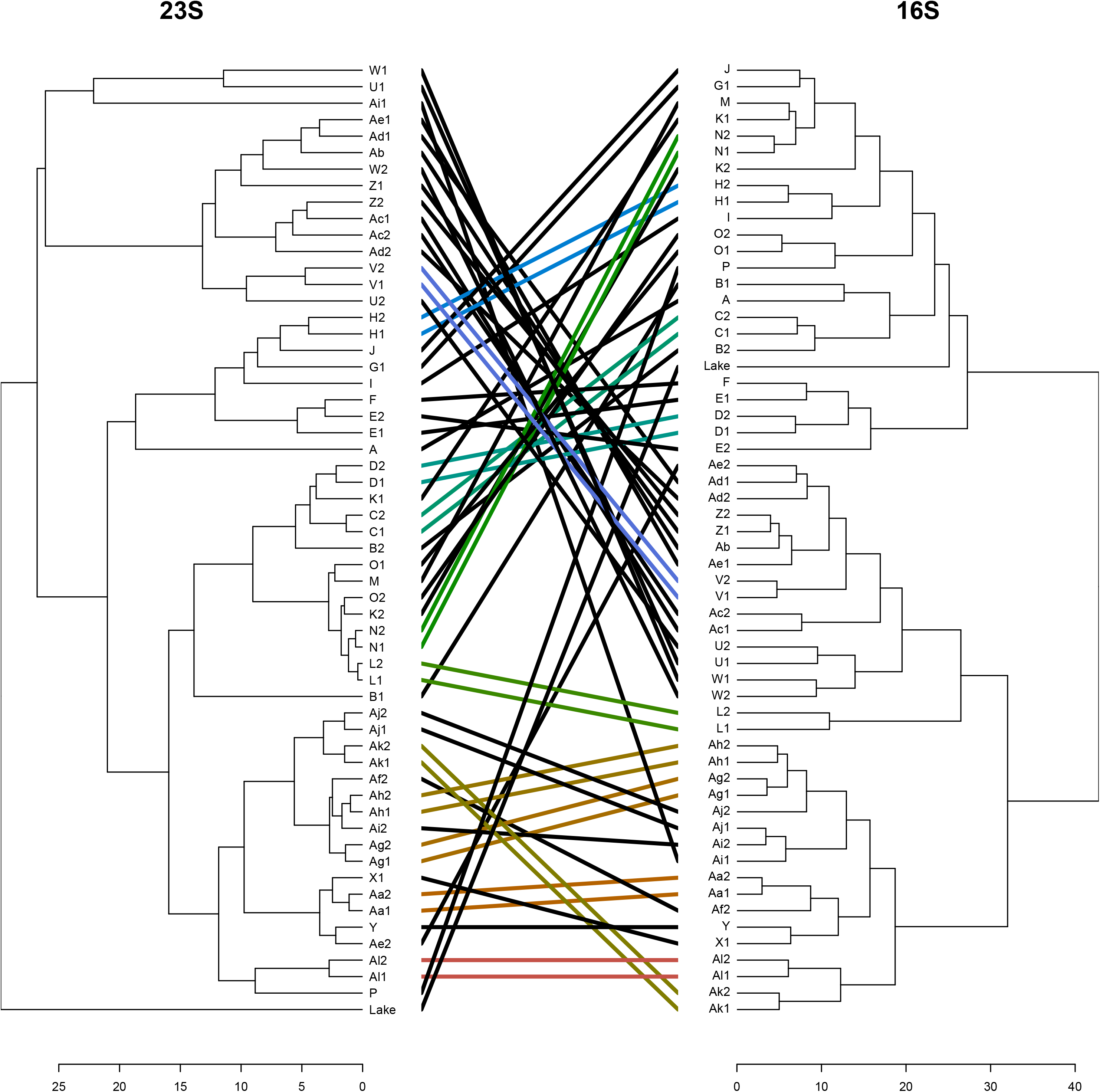
Tanglegram for the association between site hierarchical clusterings based on 23S and 16S OTU abundance. Colored lines between dendrogram tips represent similar relative placement of sites within the clustering diagram.

Compared to phytoplankton, using the same forward stepwise regression technique—top 50 OTUs, *P* < 0.01 for entry—bacterial OTUs typically explained a greater proportion of nutrient concentrations in the water. For [NH_4_ ^+^], five bacterial OTUs predicted 69% of the variation in [NH_4_^+^] compared to 38% of the variation with phytoplankton (Table 3). Sites with greater abundances of three OTUs (a Firmicutes, a Bacteroidetes, and a Proteobacteria) had higher [NH_4_^+^] while sites with greater abundances of two OTUs (an Actinobacteria and a Bacteriodetes) had lower [NH_4_^+^] (Table 3). For [NO_3_^-^], phytoplankton had predicted 61% of the variation, but the abundances of six bacterial OTUs explained 80%. [NO_3_^-^] concentrations increased with increasing abundances of two bacterial OTUs (an Actinobacteria and a Bacteriodetes) and decreased with increasing abundances of four bacterial OTUs (an Actinobacteria, a Bacteriodetes, and two Proteobacteria) (Table 3). For [PO_4_^3-^], bacteria predicted 81% of the variation in concentrations, compared to 80% for phytoplankton. [PO_4_^3-^] were lower with increasing abundances of three bacterial OTUs (an Actinobacteria, a Bacteriodetes, and a Proteobacteria) and increased with increasing abundances of a Planctomycetes OTU (Table 3).

**Table 3.** 16S OTU predictors of nutrient concentrations including sums of squares, estimates (μg L^-1^), and *P* values. Coefficients of variation for [NH_4_^+^], [NO_3_^-^], and [PO_4_ ^3-^] were r^2^ = 0.69, 0.78, 0.81, respectively.

| Nutrient | Variable | %SS | Estimate | $P$ value |
| --- | --- | --- | --- | --- |
| $[\text{NH}_4^+]$ | Intercept | | $0.2 \pm 1.3$ | 0.84 |
| | OTU13 | 21.2% | $167.8 \pm 36.7$ | <.001 |
| | OTU17 | 28.8% | $-133.1 \pm 25$ | <.001 |
| | OTU15 | 9.2% | $-52.5 \pm 17.4$ | 0.004 |
| | OTU25 | 8.2% | $138.0 \pm 48.4$ | 0.006 |
| | OTU45 | 32.6% | $899.3 \pm 158.6$ | <.001 |
| $[\text{NO}_3^-]$ | Intercept | | $1875.2 \pm 125.5$ | <.001 |
| | OTU3 | 11.1% | $6966.0 \pm 1961.3$ | <0.001 |
| | OTU7 | 11.7% | $-14045.1 \pm 3850.6$ | <0.001 |
| | OTU19 | 9.6% | $-49899.6 \pm 15098.3$ | 0.002 |
| | OTU44 | 18.5% | $-37316.1 \pm 8125.9$ | <0.001 |
| | OTU25 | 31.0% | $32079.2 \pm 5393.8$ | <0.001 |
| | OTU35 | 18.2% | $-20507.3 \pm 4508.4$ | <0.001 |
| $[\text{PO}_4^{3-}]$ | Intercept | | $102.5 \pm 8.7$ | <0.001 |
| | OTU15 | 55.0% | $-916.7 \pm 105.6$ | <0.001 |
| | OTU23 | 7.8% | $-1054.3 \pm 322.7$ | 0.002 |
| | OTU30 | 11.7% | $3924.4 \pm 981.9$ | <0.001 |
| | OTU49 | 25.5% | $-3925.7 \pm 663.6$ | <0.001 |

## Discussion

Overall, this research demonstrates the potential of sequencing the 23S region in water samples to reconstruct a broad diversity of the phytoplankton assemblage and provide information on underlying environmental conditions. Here, we saw that 23S-derived phytoplankton assemblages shifted along the length of the river, paralleled shifts in bacterial assemblages, and could predict abiotic conditions such as aquatic inorganic nutrient concentrations. These results support the further development of 23S sequencing of aquatic eDNA to reconstruct phytoplankton assemblages in order to infer environmental conditions.

The patterns in phytoplankton eDNA abundance observed for the Mississippi River were similar to those in other rivers. For example, Cannon et al. sequenced both 16S and 23S along the length of the Cuyahoga River in northern Ohio, USA (Cannon et al. 2017). As with the Mississippi, in the Cuyahoga, phytoplankton 23S OTUs were spatially patterned and many phytoplankton and bacteria were correlated along the length of the river, potentially reflecting underlying environmental conditions. In another study, Craine et al. sequenced 4 primer pairs including 16S and 23S to reconstruct biotic assemblages along 475 km of the Potomac River in Maryland, USA (Craine et al. in review). As with the Mississippi River, phytoplankton assemblages were distinctly patterned along the river and were strongly associated with river size and aquatic phosphorus concentrations. For the Potomac, phytoplankton richness increased downstream, just as with the Mississippi. Among these three studies, there were strong differences in phytoplankton assemblages. For example, compared to the Mississippi River, the Cuyahoga River had a greater dominance of Cryptophytes. Although both the lower Potomac and Mississippi were dominated by diatoms, different diatoms dominated the two rivers.

Traditional sampling of large rivers with visual quantification of phytoplankton also showed many similar patterns as we observed here. For example, in the River Loire, large portions of the river were dominated by diatoms and many taxa were associated with eutrophic conditions (Abonyi et al. 2012). In the Upper Missouri/Mississippi/Ohio River basin in 2004/5, phytoplankton diatom assemblages responded to agricultural disturbance, urbanization, and eutrophication (Kireta et al. 2012b). Compared to the other two rivers, the upper Mississippi River was distinguished by its high levels of eutrophication, with many of the taxa observed here in high abundance (or congeners) being indicative of eutrophic and/or high agricultural or urban disturbance (Kireta et al. 2012b). The lower Mississippi River is also considered generally eutrophic and many of the taxa that indicated eutrophic or saline conditions were similar to those that dominated assemblages here (Bellinger et al. 2013).

Empirically, given the greater abundance of phytoplankton in waters than, for example, insects or fish, there has been greater success sequencing the eDNA of phytoplankton than larger organisms. This further favors developing the use of phytoplankton over other taxa. The ability in this study of the relative abundance of phytoplankton to predict aquatic nutrient concentrations should encourage future research to develop this technique for bioassessment. For example, sections of the Mississippi River with high abundances of *Melosira tropica* and *Navicula salinicola* or low abundances of a *Cyclotella* OTU had high [NO_3_ ^-^]. If these relationships were to hold up across different river systems and seasons, then the abundances of these species as determined by sequencing eDNA could broadly serve as an indicator of [NO_3_^-^] without having to measure it directly. Given that using a similar technique, strong relationships between aquatic nutrient concentrations and phytoplankton abundances were seen in the Potomac River, too (Craine et al. in review), this method continues to show promise as a bioassessment tool. To our knowledge, there is no theory to explain why phytoplankton diversity increases with distance downstream and/or with increased [PO_4_ ^3-^], which was also observed in the Potomac. In fact, such observations actually run counter to the popular River Continuum Concept, which postulated that after an initial increase in headwaters, diversity should decrease with increasing river size (Vannote et al. 1980). However our observed trend of increasing diversity is consistent with other measurements of Mississippi River microbial assemblages (Payne et al. 2017)(Henson et al. in review). Theory aside, it will take much larger datasets to assess whether phytoplankton diversity, in and of itself, is diagnostic for any environmental conditions.

Greater taxonomic resolution is likely possible with other primer pairs in conjunction with 23S, but there is no evidence yet that this is necessary. That said, there are still areas where more research is required before metabarcoding with 23S for phytoplankton assemblages can be operationalized. For example, the number of sequences known from diatom taxa is small fraction of the several thousand species described from North America (Kociolek 2006). Of the nearly one thousand taxa listed in the Diatoms of the US web flora (Spaulding et al. 2010), approximately one hundred have associated sequences that are currently available in GenBank. Other studies (Visco et al. 2015) report that only 28% of taxa identified by microscopy had corresponding reads in sequence data. Consequently, for diatoms, the OTUs were mapped to taxa that were most similar, with the outcome that species that are well-characterized in gene sequences (i.e. *Melosira tropica*) that have not reported from inland waters in river surveys (U.S. Geological Survey BioData).

It is possible that the OTU matches to *Melosira tropica* could reflect the presence of the very common *M. varians. Melosira tropica* has not been reported from inland waters, but *M. varians* is one of the very common river species (Potapova and Charles 2007). Although it is not expected to find *Thalassiosira rotula* in the more northern reaches of the Mississippi River, others (Visco et al. 2015) report that the common *Stephanodiscus minutulus* was included in a well-supported clade with a number of *Thalassiosira* species, at least based on the particular region examined. *Skeletonema potamos* and *S. costatum* are commonly reported from rivers with high conductivity, resulting from agricultural input (Potapova and Charles 2007). For example, both of these species have been found in national surveys in rivers including the Milwaukee River at Milwaukee WI and the Maumee River at Waterville OH.

Beyond improving reference databases, autecological information for many phytoplankton taxa exist (Reynolds et al. 2002, Padisák et al. 2009), but indices generated with 23S will need to continue to be calibrated against environmental conditions with multiple reference sites to ensure that there are not covariates driving the relationship. For example, nutrient concentrations, distance downstream, and time of sampling were all associated in this study and these other factors could be influencing the relationships we observed between phytoplankton assemblages and nutrient availability. Multiple large rivers of different nutrient status will need to be included to partition out the direct effects of nutrient concentrations from other covariates driving assemblage composition. Reference databases will also need to be expanded by sequencing a larger diversity of phytoplankton organisms and identifying taxa associated with sequences. Although eDNA-based bioassessment can occur independent of taxonomy (Apotheloz-Perret-Gentil et al. 2017), more robust, stable indices will likely require ecological information about individual taxa, too. Given the breadth of taxa sequenced with 23S, this means broad biodiversity surveys are required for all phytoplankton taxa rather than a single taxonomic group, such as cyanobacteria. Although this technique should work with periphyton also, future work should continue to test whether phytoplankton or periphyton are best for bioassessment of given environmental conditions (Kireta et al. 2012a), although previous work with traditional techniques appears to favor the utility of phytoplankton over periphyton for assessing environmental conditions in some large rivers (Reavie et al. 2010), though not others (Bellinger et al. 2013).

## Acknowledgments

This work was supported in part by the Department of Biological Sciences, College of Science, and the Office of Research and Economic Development at Louisiana State University; and the College of the Environment at the University of Washington. The authors appreciate comments from Kristy Deiner and Ian Bishop, Meredith Tyree and Nicholas Schulte for discussions on data interpretation. Sequence data are available at www.ncbi.nlm.nih.gov/Traces/study/?acc=SRP132323.

